# The immunosuppressant tacrolimus (FK506) inhibits *C. glabrata* Cdr1 efflux pump function by stabilizing the inward-facing conformation

**DOI:** 10.64898/2026.07.08.737148

**Authors:** Baccouch Rim, Benefice Theo, Zarkadas Elefterios, Samrouth Nour, Pata Jorgaq, Magnard Sandrine, Di Meo Florent, Terreux Raphael, Aguero Stephanie, Boumendjel Ahcene, Schoehn Guy, Lamping Erwin, Falson Pierre, Chaptal Vincent

## Abstract

The pathogenic yeast *Candida glabrata* is intrinsically resistant to azole antifungals through the overexpression of the multidrug transporter Cdr1. CgCdr1 detoxifies the yeast by expelling azoles out of the cell, thereby decreasing their intracellular concentration. Tacrolimus (FK506), one of the most widely used immunosuppressant medications used world-wide, has been identified as a broad-spectrum inhibitor of Cdr1 homologs in several *Candida* species. However, its mechanism of action remains unknown. We solved the cryoEM structure of CgCdr1 in complex with FK506, with or without ATP. The structure revealed that FK506 binds within the drug-binding site of CgCdr1, occupying the space occupied by Itraconazole. The hydrophobic face of FK506 stacks against the TMD1 and forms hydrogen bonds with TMD2, stabilizing a different conformation from the one adopted in FK-binding-proteins. FK506 binding triggered structural rearrangements bringing the nucleotide-binding-domains closer to the trans-membrane-domains, while stabilizing the inward-facing conformation. While ATP can still bind to the catalytic nucleotide-binding site, FK506 prevents the conformational transition required for ATP hydrolysis, thereby effectively blocking azole transport. Inter-particle variability analysis (3DVA) revealed significant conformational flexibility of FK506 within the binding pocket, with minimal transporter mobility. It allowed to visualize the conformational space occupied by the inhibitor within its binding-pocket, serving as a useful tool for inhibitor rational design. Overall, these findings demonstrate that FK506’s inhibition extends beyond competitive binding, involving allosteric modulation of the ATPase cycle.

**Significance statement:** The pathogenic yeast *Candida glabrata* exhibits intrinsic resistance to azole antifungals via the multidrug transporter Cdr1, which expels azoles from the cell. Tacrolimus (FK506), a widely used immunosuppressant, inhibits Cdr1 homologs across *Candida* species, yet its mechanism remained unknown. Here, we resolved the cryoEM structure of *Cg*Cdr1 in complex with FK506, revealing that FK506 binds to the drug-binding site, like itraconazole. Its hydrophobic face interacts with TMD1, while hydrogen bonds form with TMD2. FK506 stabilizes the inward-facing conformation, preventing ATP hydrolysis despite ATP binding, thereby blocking azole transport. Variability analysis highlighted FK506’s conformational flexibility within the pocket, offering insights for rational inhibitor design. These findings demonstrate that FK506’s inhibition involves both competitive binding and allosteric modulation of the ATPase cycle.

## Introduction

The opportunistic fungal pathogen *Candida glabrata* exhibits intrinsic resistance to azole antifungals, primarily through the overexpression of the membrane efflux transporter Cdr1 (*1–5*). By actively expelling azoles from the cell, Cdr1 reduces their intracellular concentration below cytotoxic levels, thereby limiting drug efficacy. This inherent resistance mechanism prevents azoles from inhibiting their drug target, lanosterol-14α-demethylase, an essential enzyme of the ergosterol biosynthesis pathway encoded by *C. glabrata ERG11*.

To overcome this challenge, extensive research has focused on developing strategies to inhibit Cdr1, with the aim of restoring azole susceptibility in *C. glabrata* cells. Efficient macrocyclic efflux pump inhibitors include macrolides such as a number of milbemycins (*6*) or FK506 (tacrolimus), as well as the depsipetides beauvericin or enniatin B (*7–9*). These inhibitors exhibit broad specificity, targeting Cdr1 homologs of multiple *Candida* species including the major opportunistic fungal pathogens *C. albicans*, *C. auris* and *C.* glabrata, recently renamed *Nakaseomyces glabratus* (*10*). These compounds also inhibit the archetypal fungal drug efflux pump, *Saccharomyces cerevisiae* Pdr5, the functional ortholog of *Cg*Cdr1 and a valuable model system for mechanistic studies. This cross-species activity not only underscores the conserved nature of these transporters but also highlights the potential for developing broad-spectrum azole adjuvants.

Enzyme inhibitors are typically classified as competitive, non-competitive or uncompetitive. Competitive inhibitors share structural similarities with enzyme substrates and compete for binding to the active site. Non-competitive inhibitors, in contrast, bind to a distal site and allosterically modulate the enzyme activity. Uncompetitive inhibitors bind exclusively to the enzyme-substrate complex and lock the enzyme in an inactive state; although less common, this mechanism has been documented for ABCG2 inhibition, a member of same Type-V family of ABC transporters (*11*).

Recently, the structure of milbemycin oxime in complex with *C. albicans* Cdr1 (*Ca*Cdr1) was resolved, revealing that it binds to the central substrate binding cavity - the same site occupied by fluconazole (FLC) - suggesting a competitive mode of inhibition (*12*). However, this proposed mechanism, while compelling, lacks information about the conformational ensemble of the transporter. Transporters are highly dynamic machines that undergo substantial structural rearrangements during the export of azoles from the cytosol. Thus, reducing the inhibitor’s mode of action solely to blocking drug binding may overlook critical aspects of its mode of action.

To deepen our understanding of this inhibition, we determined the structure of *C. glabrata* Cdr1 (*Cg*Cdr1) in complex with FK506, both in the absence and presence of ATP. FK506 is an effective *Cg*Cdr1 eflux pump inhibitor (*13*) and one of the most widely used immunosuppressant medications used world-wide (*14, 15*). By comparing these two conformations, we elucidated how FK506 inhibits the ATPase activity of *Cg*Cdr1. Through 3D-variability analysis (3DVA) of the cryoEM data, we visualized the structural adaptations of the protein in response to the inhibitor and characterized the conformation ensemble adopted by FK506. This approach provided a unique and detailed depiction of the inhibition mechanism.

## Results

### FK506 inhibits Cdr1 by stabilizing the inward-facing conformation

FK506 inhibits the ATPase activity of *Cg*Cdr1 embedded in native membranes with a Ki of 15 ± 0.3 nM and a Ki of 0.14 ± 0.01 µM for the purified protein (Figure 1A), consistent with previously reported values for this transporter (*16*). The difference in inhibition constants arises from the hydrophobicity of FK506, which partitions predominantly into the membrane bilayer, thereby increasing its local concentration in the native bilayer. Notably, FK506 completely inhibited the ATPase activity of the purified protein (Figure 1A), indicating that it blocks the entire transporter pool in an ATPase-inactive state.

**Figure 1:**
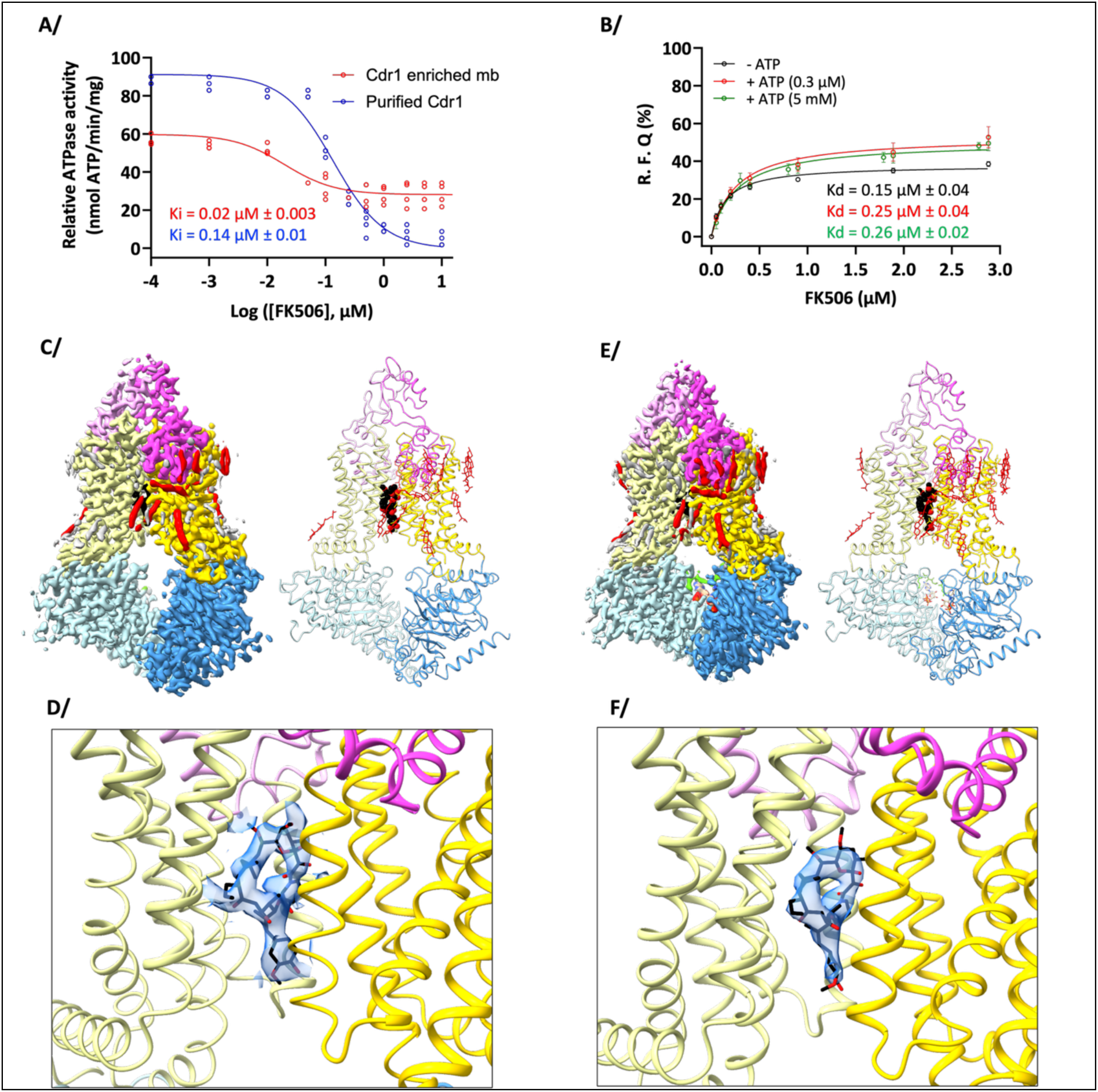
FK506 inhibition and structure in complex with *C. glabrata* Cdr1. **A/** FK506 inhibition of the ATPase activity of *Cg*Cdr1 embedded in native membranes (red) and of purified *Cg*Cdr1 (blue). **B/** FK506 binding to purified *Cg*Cdr1 apo (black) or to *Cg*Cdr1 in the presence of low (0.3 µM; red) or high (5 mM; green) ATP concentrations. R.F.Q. Relative Fluoresence Quenching. **C/** CryoEM structure of CgCdr1[sFK506|nc0|c0] in complex with FK506. The protein is colored by domains with NBD1 in cyan, TMD1 in sand, ECD1 and ECD2 in pink and purple, NBD2 in blue and TMD2 in yellow. Ergosterol molecules are colored in red and FK506 is colored by atom type with black carbons. The Coulomb map contour level is 0.379. The image to the right displays the protein as a cartoon model with FK506 displayed as spheres using the same colorings. **D/** Zoom on the FK506 binding site with the density for FK506 as transparent blue surface, contoured at 0.224, displaying the unsharpened map. The Q-score value for FK506 w*as* 0.56. **E/** Same display as in **C/** but for CgCdr1[sFK506|ncATP|cATP]. The Coulomb map contour level is 0.379. **F/** Same display as in **D/** but for CgCdr1[sFK506|ncATP|cATP] with a Coulomb map contour level of 0.0854 for the sharpened map. The Q-score value for FK506 was 0.55.

To elucidate the mechanism of FK506 inhibition, we solved the cryoEM structures of *Cg*Cdr1 in two distinct states: (1) in complex with FK506 alone (Supp-Figure 1,3), hereafter referred to as CgCdr1[sFK506|nc0|c0] where [sX|ncX|cX] denotes the occupancy of the substrate-binding site and the non-catalytic (nc) and catalytic (c) nucleotide-binding sites (NBS), respectively; and (2) in complex with FK506 and ATP-Mg2+ (CgCdr1[sFK506|ncATP|cATP]) (Supp-Figures 2-3). In the latter structure, ATP is bound at the non-catalytic NBS (ncNBS), while another ATP molecule, also without magnesium, is bound at the catalytic NBS (cNBS), clearly visible in the high-resolution reconstruction presented in Supp-Figure 4 (local resolution = 2.5-2.9 Å). Both cryoEM structures of *Cg*Cdr1 in complex with FK506 were in the inward facing (IF) conformation (Figure 1C,E), with clear densities for FK506 in the drug-binding cavity (Figure 1D,F). Like in previous structures of *Cg*Cdr1 (*16*), 13-18 ergosterol molecules decorated the membrane insertion region of *Cg*Cdr1, primarily at the catalytic face of the transporter. FK506 binding induced some conformational changes compared to the apo- or ATP-bound-states, which are discussed in detail below.

**Figure 2:**
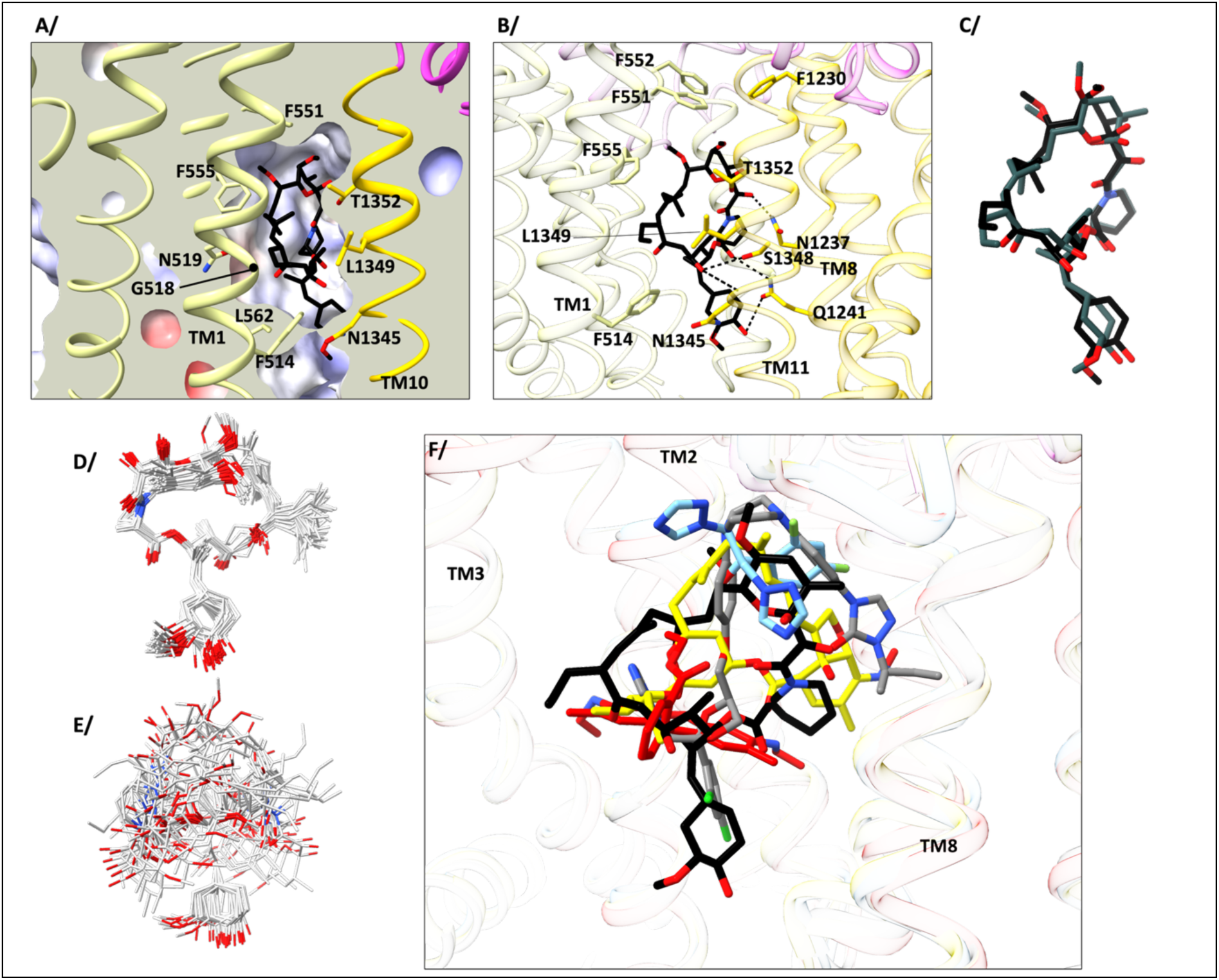
FK506 binding site. **A/** Slice through a Coulombic potential map showing the hydrophobicity of the FK506 binding site in the TMD1 of CgCdr1[sFK506|nc0|c0]. TM helices are show in cartoon colored as in Figure 1. Residues lining the drug-binding cavity are displayed as sticks and labelled. FK506 is displayed as sticks colored by atom type with black carbons. **B/** Same view as in **A/** but with hydrogen bonds between FK506 and the protein highlighted as black dashed lines. *Cg*Cdr1 is displayed as a transparent cartoon. **C/** Overlay of FK506 in *Cg*Cdr1[sFK506|nc0|c0] (black carbons) and in CgCdr1[sFK506|ncATP|cATP] complexed with ATP (grey carbons). **D/** Overlay of all 30 possible FK506 structures deposited in the PDB. **E/** Overlay of representative 20 snapshots obtained from MD simulations of FK506 in water. **F/** Overlay of CgCdr1[sFK506|nc0|c0] in complex with FK506 (black carbons) with previously published structures of *Cg*Cdr1 in complex with ITC (grey carbons), *Ca*Cdr1 in complex with FLC (cyan carbons) or MIL (yellow carbons), and *Sc*Pdr5 in complex with R6G (red carbons).

**Figure 3:**
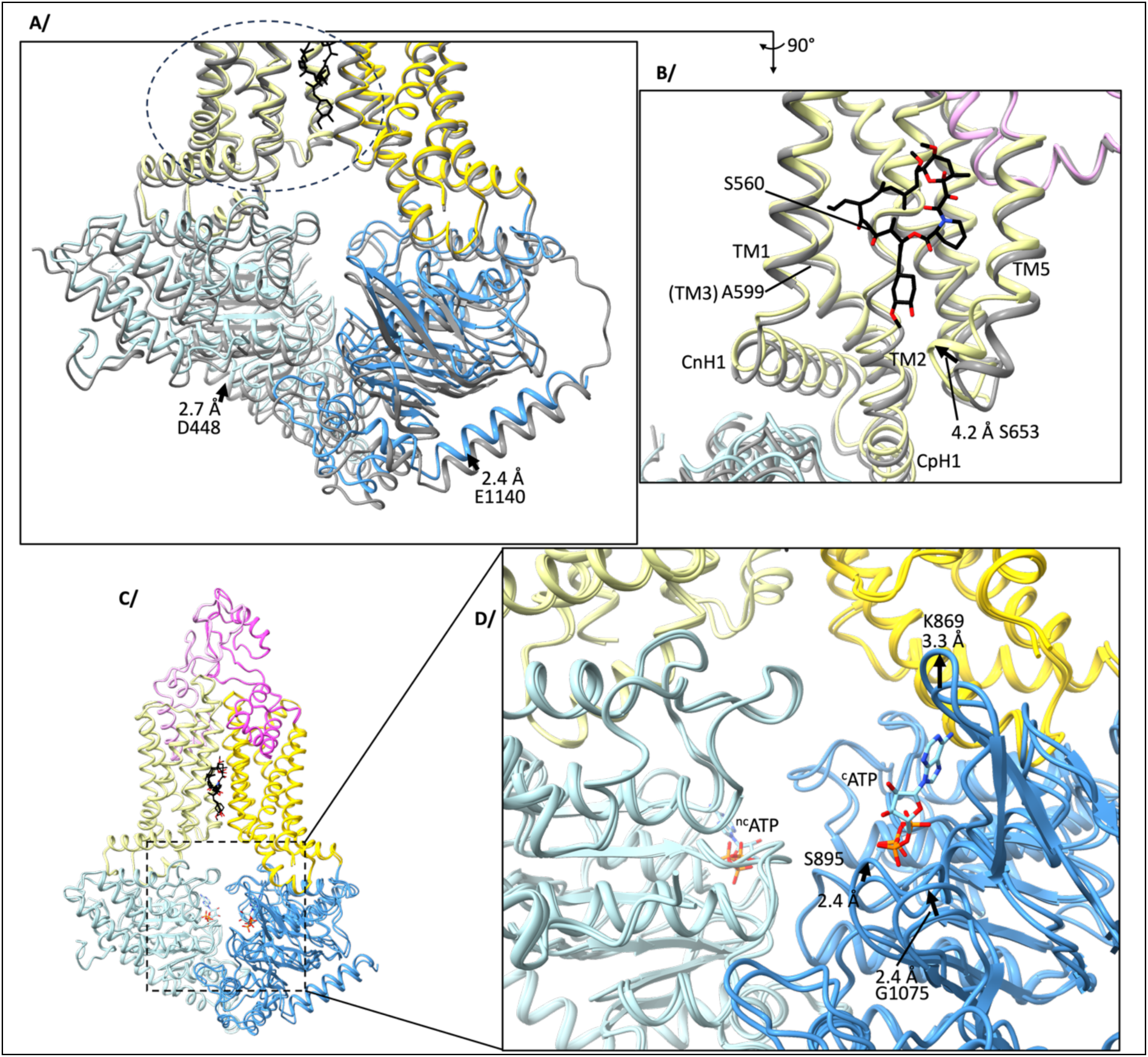
Conformational changes of *Cg*CDR1 in response to FK506 binding. **A/** Overlay of *Cg*Cdr1[sFK506|nc0|c0] colored by domain type as in Figure 1C with apo *Cg*Cdr1 (grey). The overlay was performed on TMD2 (residues 1170-1490). **B/** Zoom in view of TMD1 and the FK506 binding site, viewed from TMD2. Relevant trans membrane helices and residues displacements *a*re shown. CnH1, connecting helix H1, CpH1, coupling helix H1 **C/** Global overlay of CgCdr1[sFK506|nc0|c0] with CgCdr1[sFK506|ncATP|cATP] gave an overall rmsd of 0.6 Å. Both proteins are colored by domains as in Figure 1C. **D/** Zoom in view of the cNBS reveals local displacements of loops that are in direct contact with ATP.

**Figure 4:**
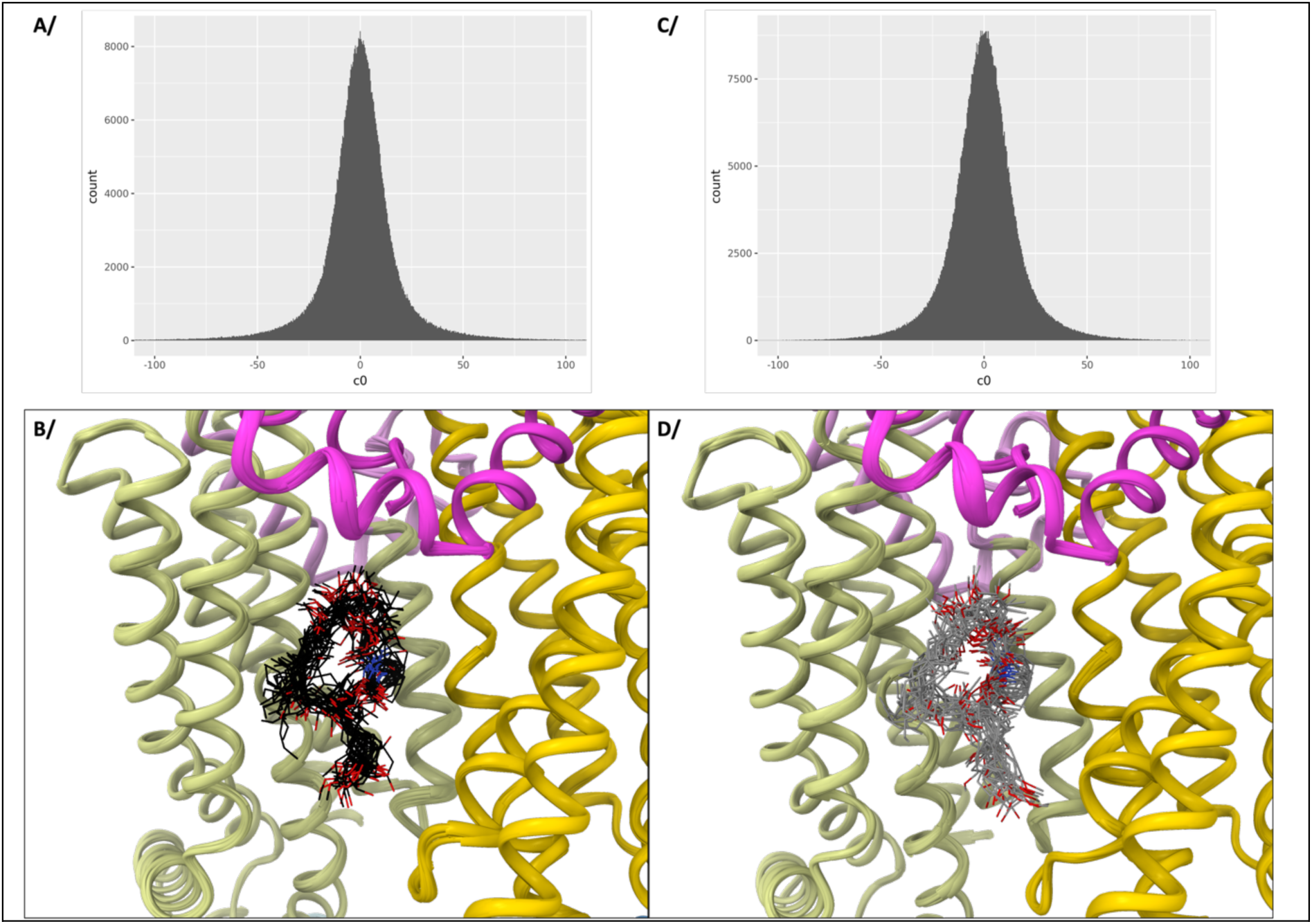
3D variability analysis of *Cg*CDR1 complexed to FK506. **A/** Particle distribution in latent space for principal component 0 as a function of latent coordinate, for CgCdr1[sFK506|nc0|c0]. **B/** Variability refinement of CgCdr1[sFK506|nc0|c0] colored as in Figure 1C, for primary component 0. C/ D/ Same as A/ and B/ but for CgCdr1[^s^FK506|^nc^ATP|^c^ATP] with FK506 colored as atom type with grey carbons.

To assess the impact of ATP-binding on FK506-binding, we initially measured the affinity of FK506 for apo *Cg*Cdr1, yielding a Kd of 0.15 ± 0.04 µM (Figure 1B). In presence of ATP, first at a 1:1 molar ratio (0.3 µM), targeting the ncNBS (where the ATP affinity is much higher) (*16*), the FK506 affinity to *Cg*Cdr1 was only slightly reduced to 0.25 ± 0.04 µM. Even at saturating ATP concentrations (5 mM, without magnesium to prevent hydrolysis), the FK506 affinity remained comparable (0.27 ± 0.02 µM) to the condition where ATP was only bound to the high affinity ncNBS. These data suggest that binding of FK506 to *Cg*Cdr1 is not affected by the presence or absence of ATP bound to one or both NBSs and it possibly stabilizes the IF conformation, locking the transporter in a state competent for ATP binding but unable to hydrolyze ATP.

### Conformational state of FK506 and FK506 binding site

A distinctive circular density with an extending tail was visible in the Coulomb potential map within the drug binding cavity of *Cg*Cdr1 in both the apo and the ATP bound structures (Figure 1D,F). This feature was present in both structures but it was absent in the original *Cg*Cdr1 apo structure (*16*), strongly suggesting the presence of an FK506 molecule occupying this space. However, fitting the ligand into this density proved unexpectedly challenging. Initial analysis of the Protein Data Bank revealed 30 structures for FK506 bound to the FK506-binding protein (FKBP), in which FK506 consistently adopted a nearly identical orientation, with only minor local adjustments (Figure 2D). Yet, this canonical conformation could not be satisfactorily fitted into the observed density, resulting in poor agreement with the experimental data and significant steric clashes with the protein. Given the circular shape of the density – consistent with the macrocyclic nature of the ligand - and the presence of a protruding tail matching the ligand’s polyketide tail, the position of the tail was unambiguous. To achieve the substantial conformational adjustments of FK506, we performed molecular dynamics simulations of FK506 to investigate its flexibility, which revealed a large conformational sampling space (Figure 2E). Some conformations were selected and manually adjusted to fit the cryoEM map leading to a good map-model correlation (Figure 1D,F). The Q-scores for the backbone structures of CgCdr1[sFK506|nc0|c0] and CgCdr1[sFK506|ncATP|cATP] were 0.71 and 0.72 and the Q-scores for FK506 in either of these two structures were 0.56 and 0.55, respectively.

FK506 binds to *Cg*Cdr1 within the drug-binding cavity, in-between TM1, TM2 and TM5 of TMD1, and TM8 and TM11 of TMD2. The TMD1 binding cavity was mostly lined with hydrophobic residues (Figure 2A) accommodating the hydrophobic face of the macrocycle. In contrast, all hydrogen bonds were formed with TMD2 residues of TM8 and TM11 – specifically N1237, Q1241, N1345, S1348 and T1352 (Figure 2B). This hydrogen-bonding network dictated the conformation adopted by FK506 within the drug-binding cavity, which differed from the conformations observed in the FKBP structures (Figure 2D). Interestingly, mutations of *Sc*Pdr5-S1360 or -T1364, equivalent to *Cg*Cdr1-S1348 and T1352, dramatically reduced the ability of FK506 to inhibit the ATPase activity of ScPdr5 and they caused altered substrate transport and transport inhibition profiles, reinforcing the importance of these TM11 residues in FK506 recognition (*17, 18*).

Structural comparison revealed that FK506 binds similarly to both the ATP-bound and ATP-free states of *Cg*Cdr1, with only modest local adjustments (Figure 2C). The roof of the FK506 binding cavity was sealed by the highly conserved aromatic residues F551, F552 of TM2 and F1230 at the top of TM8, while the front of the cavity was closed by F514 of TM1 and N1345 of TM11 acting like “saloon-doors” - a mechanism previously described for the *S. cerevisiae* Pdr5 ortholog (*19*). This arrangement likely serves as the entryway to the drug-binding cavity. Interestingly, mutations of *Sc*Pdr5-S1360 or -T1364, equivalent to *Cg*Cdr1-S1348 and T1352, dramatically reduced the ability of FK506 to inhibit the ATPase activity of ScPdr5 and they caused altered substrate transport and transport inhibition profiles, reinforcing the importance of these TM11 residues in FK506 recognition (*20*). This conserved glycine residue probably provides flexibility to TM1 and opens enough space between TM1 and TM11 for various size substrates and/or inhibitors to enter the transporter.

Interestingly, two ergosterol molecules were positioned right in front of this entrance cavity (Figure 1C,E), as already previously noted (*16*); while their role is not clear, their consistent presence at this location suggests a role in substrate recognition.

The FK506 binding site overlaps with the binding sites of itraconazole (ITC), FLC, milbemycin (MIL), and rhodamine 6G (R6G) in *Cg*Cdr1 (*16*), *Ca*Cdr1 (*12*), and *Sc*Pdr5 (*19*), respectively (Figure 2F). Structural overlay demonstrates the versatility of the drug-binding cavity in accommodating a diverse range of ligand conformations. These findings support the hypothesis that FK506 acts as a competitive eflux pump inhibitor, blocking azole binding and transport by *Cg*Cdr1, as initially proposed for *Ca*Cdr1 (*12*). Beyond acting as a competitive inhibitor, FK506 also stabilizes the conformation of *Cg*Cdr1, as discussed further below.

### The conformational changes induced by FK506 binding are similar to those induced by substrate binding

The most pronounced conformational changes were observed for CgCdr1[sFK506|nc0|c0] in the absence of ATP. When compared to the apo structure (*16*), individual domains aligned closely, with root-mean-square deviations (RMSDs) ranging from 0.6 to 0.8 Å for the entire domain. However, a global overlay of the entire structure reveals significant discrepancies (Figure 3A), highlighting an upward displacement of both NBDs toward the TMDs, with a 2 Å translation. This shift was absorbed by kinks in TM2 (S560) and TM3 (A599) of TMD1 (Figure 3B). Above these residues, the overlay with TMD1 aligned perfectly again, indicating that these kinks enable localized deformation of certain TM helices. Additionally, FK506 binding induced a local rearrangement at the junction of TM4 and TM5. These helices shifted towards the ligand closing the drug-binding cavity in an induced fit type mechanism. S653 of TM5 underwent a substantial 4.2 Å displacement toward the polyketide tail of FK506, facilitating ligand interaction.

Interestingly, binding of FK506 and ATP to CgCdr1[sFK506|ncATP|cATP] resulted in a nearly identical structure as the structure complexed with ITC (rmsd of 0.6 Å over the entire structure). CgCdr1[sFK506|ncATP|cATP] was also structurally very similar to CgCdr1[sFK506|nc0|c0] with almost all parts of the structure overlapping perfectly (rmsd also 0.6 Å over the entire structure), and local adjustments in ATP-binding loops were observed at the cNBS (Figure 3C,D).

These movements suggest that in the apo state, the NBDs are resting further apart from the TMDs than in the ATP-bound CgCdr1[sFK506|ncATP|cATP] state. It was initially thought that binding of ATP at the ncNBS induces an ATP-bound competent conformation at the cNBS that enables subsequent ATP binding and hydrolysis, but it would appear that substrate alone (i.e. FK506) can induce a similar conformational change. This conformational change probably originates from local deformations of the substrate-binding site via hydrophobic interactions with TMD1. The upward movement of the connecting helix (CnH1) attached to the N-terminus of TM1 and the coupling helix (CpH1) between TM2 and TM3 possibly induce an upward shift of NBD1 bringing it closer to the TMD region of the transporter. This movement might also induce NBD2 to shift upwards and prime *Cg*Cdr1 for ATP-binding to the cNBS. It would seem that FK506 stabilizes an ATP-bound IF conformation, not competent for ATP hydrolysis as the cNBS does not form a competent conformation for hydrolysis; the FK506-bound structures differ clearly from the vanadate-trapped state reported previously (*16*).

### Conformational 3D variability of FK506 within the drug-binding cavity

CryoEM reconstructions are derived from a set of particles representing the 3D volume in all viewing orientations, as they are frozen in vitreous ice. A corollary of this assumption is that each particle differs a little from the next depending on the structural heterogeneity of the sample at the chosen experimental conditions. In this study, we analyzed two cryoEM reconstructions of *Cg*Cdr1 in complex with FK506 to investigate the structural variability associated with inhibitor binding. We applied 3DVA (Cryosparc) (*21*) to quantify the variance for each voxel in the particle datasets, generating a series of maps that depict the conformational variability of the dataset. Using *phenix.varref* (*22*) we then produced models spanning the entire range of observed conformations. This approach has been used successfully in previous studies to explore the conformational space of the ABC transporter BmrA in the absence or presence of ligands (*23*), to assess the effects of single-site mutations on the asparagine synthase (*24*), and to reveal a piston-like motion of the ABC signature motif or the C-helix that initiates a cascade of conformational changes following ATP hydrolysis of *Cg*Cdr1 (*16*). Here, the quality of the reconstructions allowed us, for the first time, to employ 3DVA to observe how FK506 adopts a number of different conformations when binding to *Cg*Cdr1 in the presence or absence of ATP. This enabled accurate modelling of the inhibitor along the full trajectory of possible conformations (Supp-Figure 5-6).

Analysis of the latent space showed a gaussian distribution of variance for the first primary components, consistent with the quality of reconstructions (Figure 4A,C). The variability analysis will thus typically reveal how different the particles are from the central reconstruction. Overall, the *Cg*Cdr1 backbone exhibited limited movement, with a maximum displacement of 1.1 Å for both CgCdr1[sFK506|nc0|c0] and CgCdr1[sFK506|ncATP|cATP] (Figure 4B,D). In stark contrast, FK506 explored a significantly larger conformational space, with atomic displacements of up to 3.3 Å observed for CgCdr1[sFK506|nc0|c0] and 4.4 Å for CgCdr1[sFK506|ncATP|cATP]. The range of possible conformations adopted by FK506 highlights significant flexibility of the inhibitor inside the drug binding pocket. This observation demonstrates how ligands can explore a rather large volume within the drug binding cavity, possibly a hallmark of the polyspecificity characteristic of multidrug transporters.

## Discussion

FK506 (tacrolimus) is a highly potent inhibitor of *Cg*Cdr1, yet its mechanism of inhibition remains unclear. To gain structural insights, we determined the cryo-EM structures of *Cg*Cdr1 in complex with FK506, both in the presence and absence of ATP. The inhibitor was unambiguously localized within the drug-binding cavity, mirroring the position of milbemycin oxime in C*a*Cdr1 (*12*). FK506 adopts a different orientation from the one bound to the FKBP. The FKBP-FK506 complex acts as a potent immunosuppressant by inhibiting the calcineurin phosphatase activity. This property grants FK506 a role in treating lupus or to prevent graft rejection of transplant patients (*25*). Complexed with *Cg*Cdr1, the hydrophobic face of FK506 interacts with TMD1 residues and leverages unwinding of TM2 to nestle into the drug binding cavity. In contrast, FK506 forms several hydrogen bonds with TMD2 residues. The conformational flexibility of FK506, supported by molecular dynamics (MD) simulations (**Figure 2E**), enables its binding to *Cg*Cdr1. Once bound, FK506 stabilizes the IF conformation of the transporter. Although ATP can still bind to the cNBS of *Cg*Cdr1, the transporter is now unable to transition the cNBS into the closed conformation required for ATP hydrolysis. Consequently, the inhibition mechanism involves not only high-affinity binding of FK506 to the drug-binding pocket but also the inhibition of ATP hydrolysis, effectively blocking active azole transport by *Cg*Cdr1. Thus, FK506’s mode of action extends beyond mere competition for the drug-binding site.

The inter-particle variability analysis revealed that FK506 explores a significant conformational space within the drug-binding cavity, while the transporter itself exhibits minimal mobility. This indicates that, although the cavity adapts to the ligand, the adaptation is relatively loose, allowing FK506 to adopt multiple poses within the drug-binding site. This observation resonates with the ability of *Cg*Cdr1 and related fungal multidrug eflux pumps to bind and transport a plethora of chemically-unrelated drugs. Previous studies have shown that multidrug resistance (MDR) transporters can accommodate various ligands in their drug-binding cavities, adopting diverse poses to bind multiple substrates. The 3DVA of FK506 is the first experimental report of ligand mobility within a drug-binding site. This type of analysis is invaluable for visualizing the behavior of inhibitors in complex with proteins and for elucidating their mechanisms of action.

An unexpected conformational change was observed for *Cg*Cdr1 in response to FK506 binding which involves priming of the ATP binding sites through a series of induced-fit local deformations. These deformations originated at the kink regions of TM2, TM3 and TM5 upon FK506 binding and propagate all the way to the NBDs. As a result of FK506 binding to apo *Cg*Cdr1 ([sFK506|nc0|c0]), the two NBDs adopted the same position as when ATP was boundto *Cg*Cdr1 ([sFK506|ncATP|cATP]). This suggests that FK506 possibly induces a conformational change similar to the one triggered by ATP binding to the ncNBS. One could hypothesize that FK506 can stimulate the ATPase activity of *Cg*Cdr1 in the presence of a substrate, as it would reduce the steps required to bring the NBDs together. However, such ATPase stimulation has not been observed for *Cg*Cdr1 or other proteins in the same family (e.g., *Sc*Pdr5). Initially, it was thought that the high affinity for the ncNBS means that ATP is constantly bound to thencNBS, priming the transporter to receive substrate and have ATP bound to the cNBS, potentially explaining the absence of substrate-induced ATPase stimulation. Alternatively, the conformational change induced by FK506 may be a rapid step with limited influence on the overall ATPase cycle, with the rate-limiting step occurring elsewhere in the reaction cycle of *Cg*Cdr1. Previously, the absence of ATPase stimulation in PDR transporters suggested that substrates have limited impact on the transporter’s conformational ensemble, unlike other ABC transporters. This study provides the first report of a ligand-binding-induced conformational change for this family of fungal PDR transporters.

## Material & Methods

### Materials

Bacto-yeast extract and bacto-peptone were purchased from Difco Laboratories, Detroit, MI, USA. Luria Bertani (LB) and Yeast-Peptone-Dextrose (YPD) media were purchased from Carl Roth Gmbh & Co. Kg (Karlsruhe, Germany). Trans-PCC-α-M (*26*) (PCC) was purchased from Glycon Biochemicals, Luckenwalde, Germany. Dicarboxylate oside detergent 9b (*27*) (DCOD 9b) was a gift from CALIXAR, Lyon, France. EDTA-free antiprotease mix was purchased from Roche SAS, Boulogne-Billancourt, France. HiFliQ Nickel-NTA columns were purchased from Generon, Slough, UK. Superdex 200 column was purchased from CYTIVA Europe Gmbh, Velizy-Villacoublay, France. Amicon®ultra-centrifugal filters were purchased from Merck KGaA, Darmstadt, Germany. Oligomycin, FK506 and itraconazole were purchased from Sigma, L’lsle-d’Abeau Chesnes, France, and prepared as stock solutions in 100% DMSO.

#### Strains

The yeast strain AD/CgCdr1-His6 used in this study for producing *Cg*Cdr1 was derived from the *Saccharomyces cerevisiæ* AD1-8u- strain in which genes coding for the ABC exporters Yor1, Snq2, Pdr5, Pdr10, Pdr11, Ycf1, Pdr15, and the transcription factor Pdr3 were deleted, rendering the strain hypersensitive to drugs (*28*). *Cg*C*DR1* (Q6FK23; Uniprot database) was derived from the *C. glabrata* laboratory strain CBS 138, and cloned with a C-terminal hexa-histidine tag integrated at the *PDR5* locus of AD1-8u^−^ to form AD/CgCdr1-His6 (*29*). AD1-8u^−^and AD/*Cg*Cdr1-His6 were a gift from Richard Cannon and Erwin Lamping.

#### Cdr1 expression and purification

Following the protocol described in (*30*), 3 L of AD/CgCdr1-His6 cells were grown in 0.5 L volumes of YPD medium [1% (w/v) yeast extract, 2% (w/v) peptone, and 2% (w/v) D-glucose] in 3-L Fernbach flasks at 30 °C with shaking at 200 rpm to ensure vigorous aeration. Logarithmic AD/CgCdr1-His6 cells were harvested by centrifugation at 7,000 × g when the culture reached an OD₆₀₀ of 4, washed once with ice cold distilled water, and resuspended in ∼150 mL ice-cold lysis buffer containing 20% (w/v) glycerol, 0.5 mM EDTA, 1 mM PMSF, 1× CLAPA protease inhibitor cocktail, and 50 mM Tris-HCl (pH 7.5). PMSF was prepared as a 200 mM stock solution in isopropanol and stored at −20 °C. The ice-cold cell suspension was lysed immediately at 1.5 kbar using a cell disruptor (Constant Cell Disruptor Systems) with a single pass at 4 °C. Cell debris were removed by centrifugation at 3,000 × g for 10 min. The clarified supernatant was subsequently diluted twice with lysis buffer and centrifuged at 20,000 × g for 1 h in a Beckman JLA 16.250 rotor. The pelleted membrane fraction (C20K) was resuspended in 10 ml ice cold 20% (w/v) glycerol, 1 mM PMSF, 1 × CLAPA antiprotease cocktail and 50 mM Tris-HCl (pH 7.5).

The total protein concentration of the crude membrane preparation was determined using the Lowry method (Pierce) using BSA as standard and stored at ∼10 mg/ml at −80 °C for subsequent ATPase activity assays or for purification. *Cg*Cdr1 extraction was carried out at a protein concentration of 5 mg/mL in the buffer described above supplemented with 1% (w/v) PCC and 0,1 % (w/v) DCOD-9b (*27*). The mixture was incubated for 1 h at 4 °C under gentle agitation. The detergent-soluble fraction was obtained by centrifugation at 20,000 × g for 30 min at 4 °C and loaded onto a 5 ml HiTrap™ IMAC Sepharose™ FF column (Cytiva) preequilibrated with 5 volumes of equilibration buffer containing 50 mM Tris Hcl (pH 7.5), 150 mM NaCl, 10 mM imidazole, 1 × CLAPA antiprotease cocktail, 0.005 % (w/v) PCC and 0.0005 % (w/v) DCOD-9b. The resin was then washed with the equilibration buffer containing 20 mM imidazole and *Cg*Cdr1 was eluted by increasing the imidazole concentration to 150 mM. The pool of about 10 mL was 10x-concentrated using Amicon ultra-centrifugal filters (Millipore) with a cutoff of 100 kDa at 1,000 x g to limit detergent over-accumulation and then loaded on a Superose 6 increase 10/300 GL column (Cytiva) with a running buffer containing 50 mM HEPES-HCl (pH 7.5), 100 mM NaCl, 0.005 % (w/v) PCC and 0.0005 % (w/v) DCOD-9b. The peak eluted at 15.5 mL corresponding to *Cg*Cdr1 was collected, pooled and either immediately used for Cryo-EM and ATPase activity assays or frozen in liquid nitrogen and stored at -80 °C.

#### FK506 binding assays

Binding of FK506 (± ATP) was carried out by probing intrinsic fluorescence quenching as a function of the FK506 concentration using a SAFAS 15 Xenius spectrofluorimeter (SAFAS, Monaco). *Cg*Cdr1 tryptophan residues and N-acetyl tryptophan amide (NATA) used as negative control were excited at 290 nm with a slit width of 5 nm and their fluorescence emission spectra were recorded between 310 and 380 nm with a 3 nm slit width. Fluorescence change was recorded at a constant photo multiplicator voltage of 740 V at 0.3 µM *Cg*Cdr1. NATA, resuspended in the same buffer as *Cg*Cdr1, was used at a concentration equivalent to that of *Cg*Cdr1 tryptophan residues (5 µM). Experiments were performed in a final volume of 200 µL in a quartz cuvette, to which increasing amounts of FK506 were added, at a controlled temperature of 25°C and incubation times of 1 min. Influence of DMSO was subtracted to the raw fluorescence and kept below 0.7 % (v/v) final.

Fluorescence emission curves were integrated between 310 and 380 nm, the effect on *Cg*Cdr1 was compared to the one of NATA using the equation f = 100*ABS((F/F0)CgCdr1-(F/F0)NATA), where F is the fluorescence emitted by *Cg*Cdr1 or NATA at a given ligand concentration and F0 being the fluorescence emitted before ligand addition. FNATA did not change in presence of FK506 while FCgCdr1 increases, resulting in a relative fluorescence increase plotted as a function of the FK506 concentration. Final plots were generated with GraphPad 10.1.2 (Prism), using the equation for ligand saturation of one site-specific binding, Y=Bmax*X/ (Kd + X) where X is the FK506 concentration, Bmax is the maximum fluoresence and Kd is fitted.

#### ATPase activity

ATP hydrolysis was measured by quantifying the released inorganic phosphate (Pi). *Cg*Cdr1-associated ATPase activity was assessed by suspending 25 µg/10 µL of the C20K membrane fraction or 5 µg *Cg*Cdr1/10 µL in a final volume of 100 µL buffer containing 8 mM MgCl₂, 5 mM ATP, and 60 mM Tris-HCl (pH 7.5), along with 1 mM NaN₃ and 5 mM KNO₃ as F₁F₀-ATPase and pyrophosphatase inhibitors. FK506 were added to the reaction mixture at concentrations ranging from nM to µM. After 30 min incubation at 30 °C, the reaction was stopped by adding 100 µL of a freshly prepared cold solution containing 10 g/L SDS, 10 g/L ammonium molybdate, 2 g/L ascorbic acid, and 40 g/L H₂SO₄. The amount of released Pi was quantified by measuring the absorbance at 880 nm using a SAFAS Xenius plate reader. A standard curve was generated using known amounts of KH₂PO₄ (10–100 nmol) prepared under the same conditions in the absence of C20K or *Cg*Cdr1. Data was analyzed using GraphPad 10.1.2, with the equation Y=Bottom + (Top-Bottom)/(1+(X/IC50)), where X is the FK506 concentration and IC50 is fitted.

#### Sample preparation for cryo-EM and grid preparation

Freshly purified CgCdr1 was concentrated to 9 g/L using Amicon®ultra-centrifugal filters with a cutoff of 100 kDa at 1,000 x g. FK506 (Sigma) was solubilized in 100% DMSO at a concentration of 4 mM. For the *Cg*Cdr1[sFK506|nc0|c0] condition, 0.9 µL of SEC buffer and 0.3 µl of FK506 solution were placed first at the bottom of a 1.5 mL microcentrifuge tube. Finally, 23.8 µL of purified CgCdr1 was added with immediate pipetting, yielding final concentrations of 8.5 mg/mL (53.3 µM) for *Cg*Cdr1 and 48 µM for FK506. 3 µL of this mixture were immediately applied to freshly glow discharged (on air for 50 s at 30 mA, Emitech Glow Discharge) UltraAufoil Au/Au 1.2/1.3 grids (Quantifoil) at 20 °C and 100 % humidity using a Vitrobot Mark IV (Thermo Fisher Scientific). Excess liquid was blotted for 4 s at blot force 0 and 0.5 s drain time before vitrification in liquid ethane. Grids were verified and collected on a Thermo Scientific Glacios Cryo-TEM at 200 kV equipped with a Falcon4i direct detector (IBS, Grenoble). For the *Cg*Cdr1[sFK506|ncATP|cATP] condition, the sample was treated identically with the only difference being the addition of 0.9 µl of 141.5 mM ATP-Mg2+ (5 mM final) to the FK506 volume prior to the addition of *Cg*Cdr1. Image acquisition parameters and analysis statistics are summarized in Supplementary Table 1.

#### Cryo-EM image processing

Both datasets were processed similarly using cryoSPARC v4.7.1 (Structura Biotechnology Inc, Toronto). Motion correction was performed with PatchMotion, and contrast transfer function (CTF) parameters were estimated from averaged movies using PatchCTF. To remove any form of bias, automatic particle selection was performed using a blob picker without the use of templates on a subset of images, followed by 2D classification. The good classes were used as a template for template picking of the whole dataset and particle extraction. Initial maps were calculated *ab initio*, followed by hetero-refinement to remove junk particles. The final maps were refined using cryoSPARC’s non-uniform refinement feature without any applied symmetry. Detailed image processing parameters can be found in Supplementary Table 1. Given the quality of the grids with large number of particles per hole and the large datasets collected, a significant number of particles were used for the reconstructions. Trials to reduce this number in anyway only resulted in worse or similar resolutions as soon as about 100k particles were included in the reconstructions. Different local refinement strategies were employed with large masks or focused masks around the ligand-binding site, which did not meaningfully improve the quality of the reconstructions. 3DVA was performed using particles from consensus refinements, using a filter resolution to 4.0 Å and a 20 Å dilated mask, displaying 3 primary components at the full resolution range of the dataset.

#### 3D-model building and refinement

The apo-structure of CgCdr1[s0|nc0|c0] (PDB 9s4t) was docked into the cryo-EM reconstruction using phenix.dockinmap (Phenix) and improved with iterative rounds of manual building in Coot and Isolde followed by real_space_refine in Phenix. The final model was built using both sharpened and unsharpened maps. The final model was validated with MolProbity and deposited in the Protein Data Bank under the accession code 32CV and in the electron microscopy database EMDB-58809. The figures were constructed using and UCSF ChimeraX (University of California, San Francisco) (*31*). For the ATP-bound CgCdr1[sFK506|ncATP|cATP] structure, the initial structure was the structure in complex with Itraconazole and ADP (PDB 9s2h). PDB and EMDB codes (32DC and EMD-58811 respectively) and model statistics for both structures are provided in Supplementary Table 1.

Fitting FK506 inside the circular map within the drug-binding site was not trivial. Using the previous orientation of FK506 in FKBP proved impossible to adjust by rigid-body adjustments, resulting in clashes with the protein. The circular shape of the map with a protruding feature matches the circular ring of FK506 with its polyketide tail. As a result, only two possible orientations were feasible (rotated by 180°), but both resulted in clashes with the protein. Attempts to alternatively fit the macrocyclic ring into the circular density, regardless of the polyketide tail’s position, also led to poor fitting of the macrocycle and multiple steric conflicts with the transmembrane (TM) helices of *Cg*Cdr1. Thus, substantial conformational adjustments of FK506 were required to achieve a viable fit within the observed map. Molecular dynamics simulations of FK506 revealed the ligand’s flexibility. Some conformations were selected to fit the cryoEM map and were manually adjusted leading to a good map-model correlation with a good stereochemistry.

#### 3D-variability analysis and variability refinement

Three-dimensional variability analysis (3DVA) (*21*) of CgCdr1 was performed in CryoSPARC v4.7. The analysis was conducted on particles and masks output from Non-Uniform refinement jobs for each dataset. The masks were previously dilated by 20 Å to extend the search space. The filter resolution was set to 4 Å, no symmetry was imposed (C1), and calculated for 3 modes. The display job was calculated in simple or intermediate modes to 3 Å resolutions, with a series of 20 maps. A series of 20 models fitting the 20 maps were generated using *phenix.varref* (*22*) against the maps and the high resolution model. ERG molecules were previously removed and loose restraints for FK506 were included to allow deformation without loosing stereochemistry. Results of *phenix.varref* outputs were analyzed using ChimeraX-1.12. Q-scores were computed using the Q-score plugin of ChimeraX.

#### Molecular dynamics simulation of FK506

To facilitate the interpretation of the cryoEM densities and account for the conformational flexibility of tacrolimus outside the context of FKBP, we performed extensive molecular dynamics (MD) simulations using the AMBER suite (*32*). Tacrolimus was modeled using the General Amber Force Field 2 (GAFF2), with parameters and partial charges derived from a previous study (*33*). To ensure exhaustive sampling of the ligand’s conformational landscape, two distinct simulation environments were employed: (i) a solvated system utilizing the OPC water model (*34*) with a 12 Å buffer and (ii) a vacuum system to maximize the exploration of high-energy barriers and diverse macrocyclic ring puckers. For each environment, fifteen independent replicas were conducted, each initiated with a unique random seed to ensure statistical robustness. The solvated systems underwent a rigorous multi-stage relaxation protocol starting with five successive minimization phases. The initial stage focused on the localization of solvent oxygen atoms and counterions while maintaining a 100 kcal.mol-1.Å-2 harmonic restraint on the ligand, followed by a second stage dedicated to the minimization of all hydrogen atoms. Three subsequent unrestrained minimization phases were performed, with the final stages extending up to 107 cycles to ensure the removal of all local steric clashes. Following minimization, the systems were gradually heated from 0 to 300 K over 500 ps in the NVT ensemble using the Langevin thermostat with a collision frequency of 2.0 ps−1. During this heating phase, temperature ramping was controlled to maintain a linear gradient over the first 300 ps before holding the system at 300 K. Equilibration was continued in the NPT ensemble at 1 atm and 300 K using isotropic position scaling and a pressure relaxation time of 2.0 ps. An initial 2 ns equilibration phase was performed with ligand restraints, followed by an unrestrained 50 ns production-level equilibration. All simulations utilized a 2 fs timestep with the SHAKE algorithm (*35*) to constrain bonds involving hydrogen atoms. Non-bonded interactions were truncated at a 10 Å cutoff, and long-range electrostatics were treated using the Particle Mesh Ewald method (*36*). Snapshots were extracted from the production trajectories across all fifteen replicas and clustered based on the root-mean-square deviation of the macrocyclic ring. Post-processing and conformational analysis were conducted using the cpptraj software (*37*). We performed conformational clustering to identify representative states of the ligand. A k-means clustering algorithm (*38*) was employed to partition the ensemble into 20 distinct structural clusters based on the distance matrix error (DME) of all non-hydrogen atoms. This metric was chosen to capture the internal conformational variance of the macrocycle independently of global rotation and translation, processing half the production frames. To facilitate accurate density fitting, the representative centroids from each cluster were then aligned onto the O31-containing macrocyclic ring of tacrolimus, as this rigid moiety provided the most unambiguous feature within the cryo-EM reconstruction map.

## Acknowledgments

We gratefully acknowledge support from the CNRS/IN2P3 Computing Center (Lyon - France) for providing computing and data-processing resources needed for this work.” A special thanks to Alexis Michon and his team for management of the IBCP computing facilities. This work used the platforms of the Grenoble Instruct-ERIC center (ISBG; UAR 3518 CNRS-CEA-UGA-EMBL) within the Grenoble Partnership for Structural Biology (PSB), supported by FRISBI and GRAL, financed within the University Grenoble Alpes graduate school (Ecoles Universitaires de Recherche). We are grateful to the Nouvelle Aquitaine regional supercomputer CALI3 (“CAlcul en Limousin”), as well as and Xavier Montagutelli (University de Limoges, France) for its computational support.

## Funding

The project was supported by the Agence Nationale de la Recherche (Française), ((French) National Research Agency): ANR-24-CE44-3527 RESTOR to VC, PF and AB and the ANR-23-CE11-0031 BmrAMeca to VC and RB, the ANR-19-CE17-0020 IMOTEP and ANR-21-CE18-0030 RAPRACLID to FDM. TB PhD was supported by the doctoral school EDISS 205. FRISBI (ANR-10-INBS-0005-02) and GRAL, financed within the University Grenoble Alpes graduate school (Ecoles Universitaires de Recherche) CBH-EUR-GS (ANR-17-EURE-0003) to GS and LZ. The IBS-ISBG EM facility is supported by the Auvergne-Rhône-Alpes Region, the Fondation Recherche Medicale (FRM), the fonds FEDER and the GIS-Infrastructures en Biologie Santé et Agronomie (IBISA). EL was supported by the Marsden Fund grant MFP-UOO2210 awarded by the Royal Society of New Zealand.

## Author contributions

RB expressed and purified CgCdr1 and performed activity and binding assays. RB prepared the protein for cryoEM experiments. LZ prepared the cryoEM grids and collected the data. VC, RB and NS processed the cryoEM data, VC did the model building. NS performed expression and purification of *Cg*Cdr1. TB performed 3DVA and *phenix.varref* analyses. FDM performed the MD simulations. All authors analyzed the data and participated in writing the manuscript. VC lead the project.

## Competing interests

Authors declare that they have no competing interests.

---

**Supp-Figure 1:**
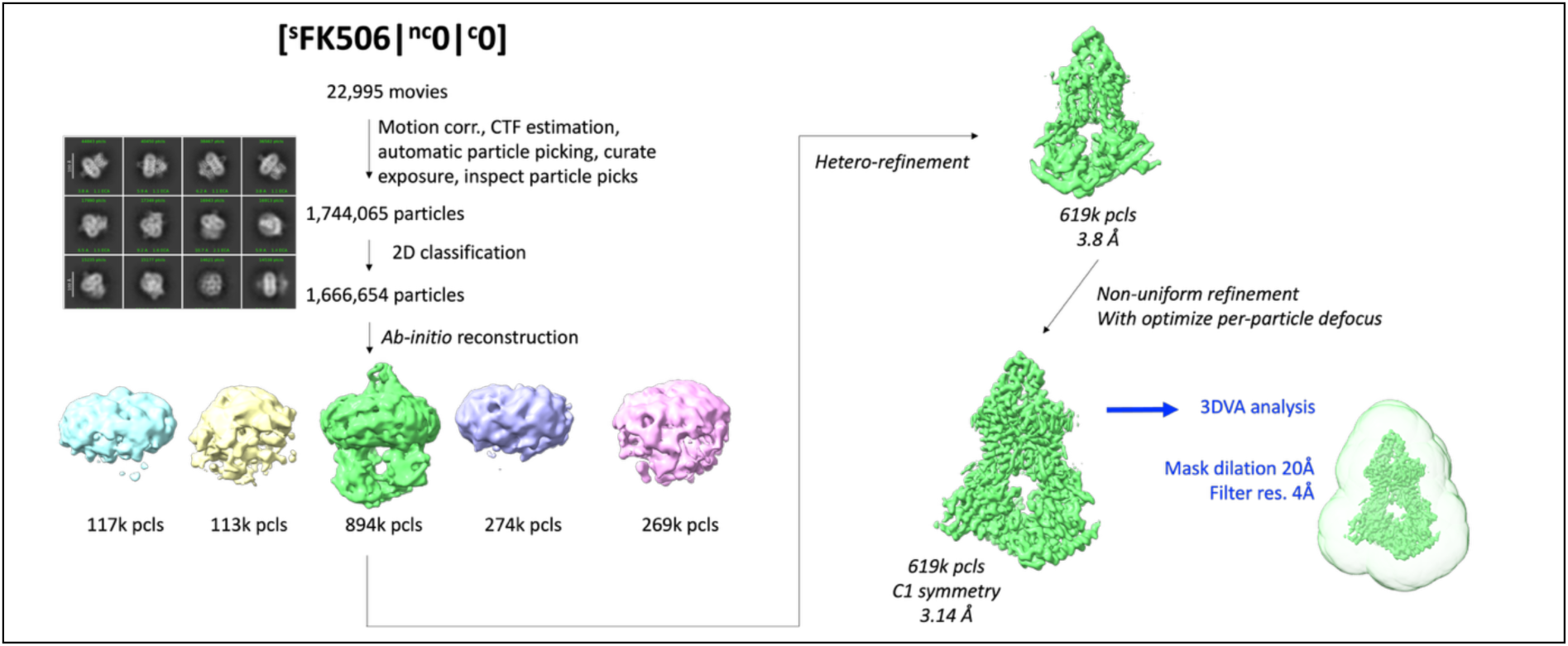
CryoEM data processing and reconstruction for *Cg*Cdr1[sFK506|nc0|c0]. Particles (pcls) are listed for each step and class, and resolutions at FSC=0.143 are listed for the latest stages of refinement. The final particle set was used for 3DVA with a mask dilation of 20 Å and a filter resolution of 4 Å.

**Supp-Figure 2:**
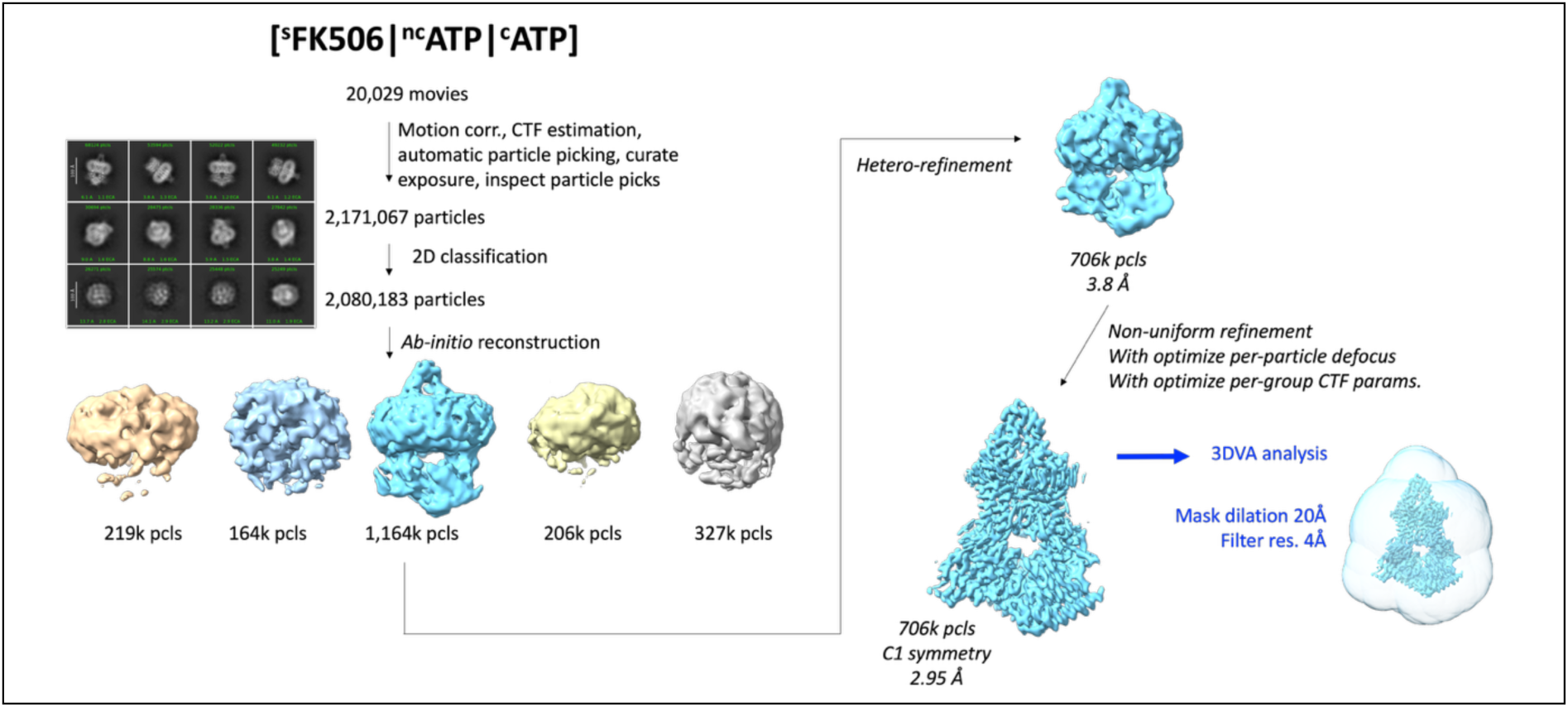
CryoEM data processing and reconstruction for *Cg*Cdr1[sFK506|ncATP|cATP]. Particles (pcls) are listed for each step and class, and resolutions at FSC=0.143 are listed for the latest stages of refinement. The final particle set was used for 3DVA with a mask dilation of 20 Å and a filter resolution of 4 Å.

**Supp-Figure 3:**
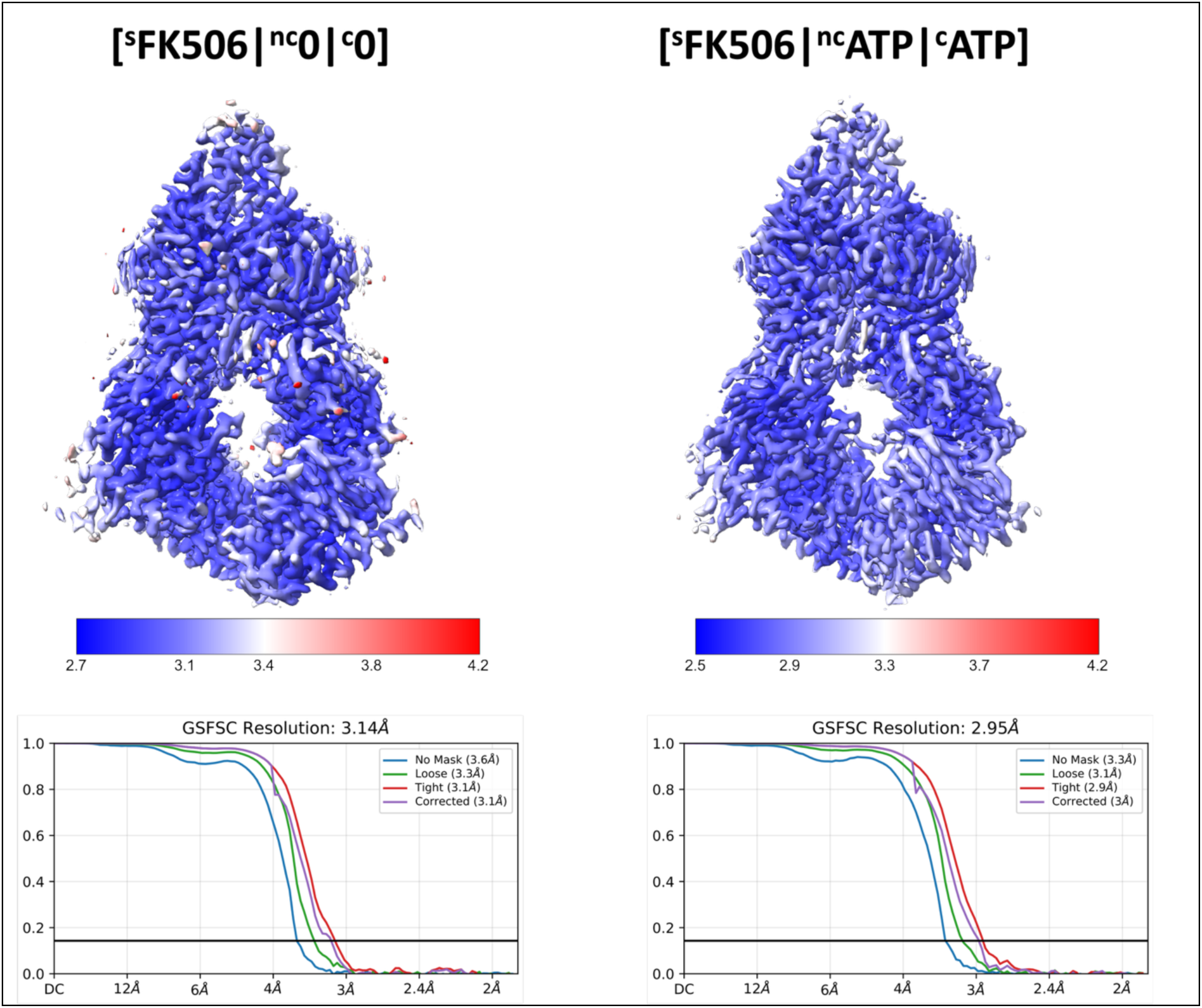
Local resolution estimations for each dataset. Top: local resolution estimates with the scale bar below. Bottom: global FSC with the global resolution estimate based on FSC0.143.

**Supp-Figure 4:**
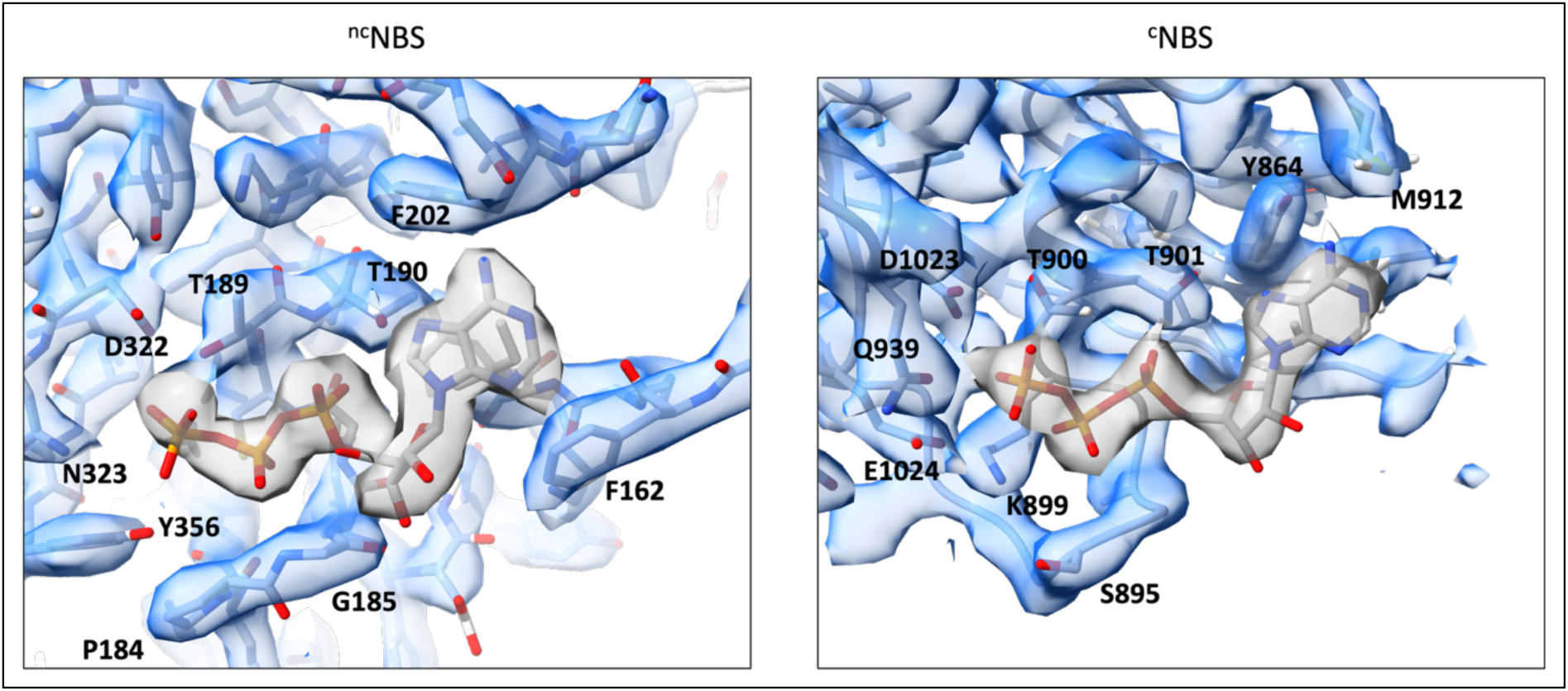
Coulomb potential maps for ATP of *Cg*Cdr1[sFK506|ncATP|cATP]. Non-catalytic NBS (left) and catalytic NBS (right) are displayed with the protein and ATP in sticks colored by atom type. The protein map is displayed in transparent blue at a map level of 0.365 for the ncNBS and 0.24 for the cNBS. The Coulomb potential maps for ATP are colored in transparent grey at map contour levels of 0.35 for the ncNBS and 0.24 for the cNBS.

**Supp-Figure 5:**
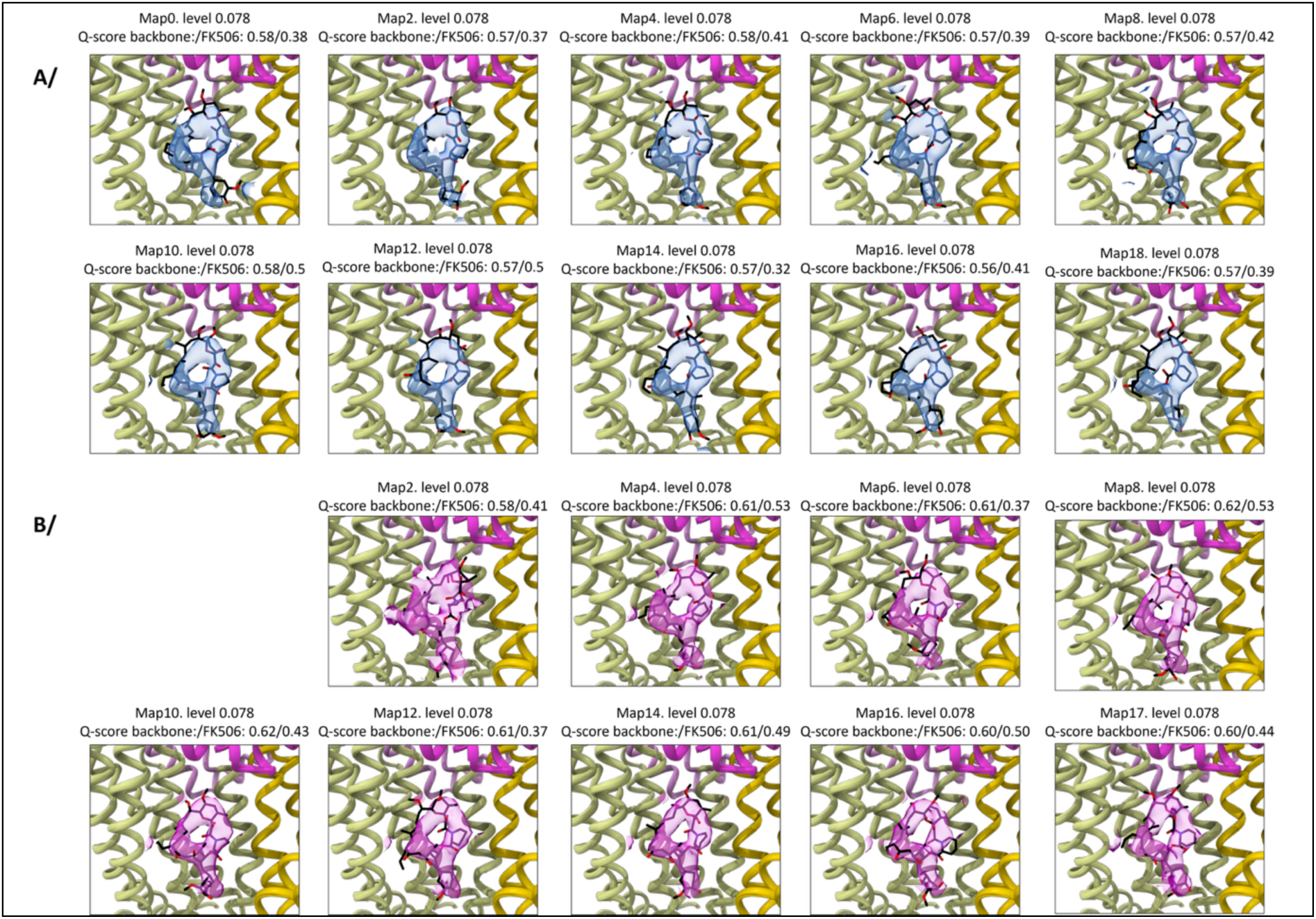
Variability analysis for *Cg*Cdr1[sFK506|nc0|c0]. **A/** FK506 binding site with the map for FK506 as transparent blue surface using the simple mode analysis (deformation of the initial volume) **B/** Same as A/ but using the intermediate mode analysis (reconstruction of new maps from subpopulation of particle) and the map for FK506 as transparent magenta surface. All maps are displayed for the component 0. Map numbers are listed above each figure with the contour level used for display, as well as the Q-score for the protein backbone and FK506. The protein displayed corresponds to the variability refinement output of each particular map.

**Supp-Figure 6:**
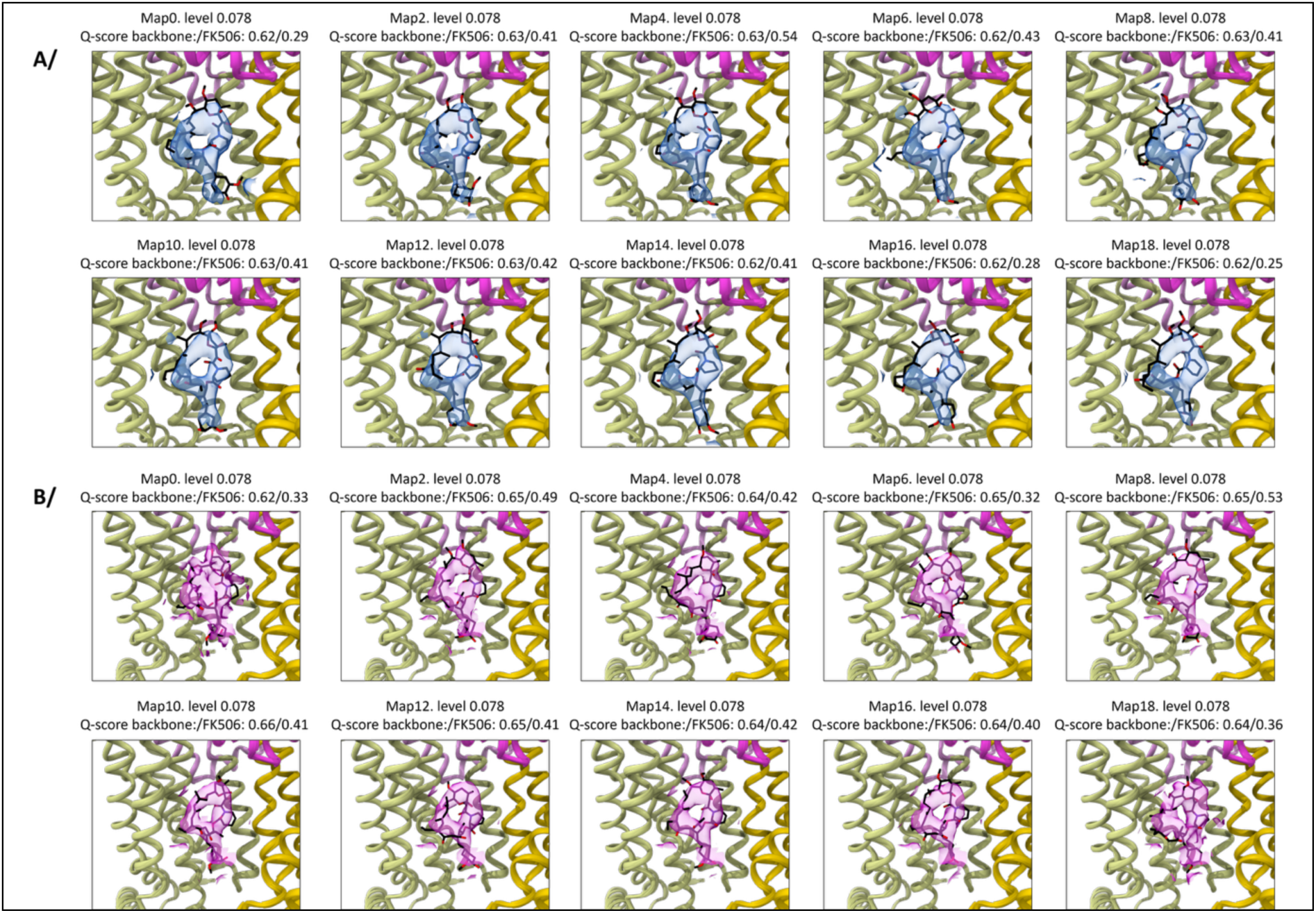
Variability analysis for *Cg*Cdr1[sFK506|ncATP|cATP]. FK506 binding site with the map for FK506 as transparent blue surface using the simple mode analysis (deformation of the initial volume) **B/** Same as A/ but using the intermediate mode analysis (reconstruction of new maps from subpopulation of particle) and the map for FK506 as transparent magenta surface. All maps are displayed for the component 0. Map numbers are listed above each figure with the contour level used for display, as well as the Q-score for the protein backbone and FK506. The protein displayed corresponds to the variability refinement output of each particular map.

|  | CgCdr1[<sup>s</sup>FK506]<sup>[nc0][<sup>c</sup>0]</sup> | CgCdr1[<sup>s</sup>FK506]<sup>[ncATP][<sup>c</sup>ATP]</sup> |
| --- | --- | --- |
| **PDB code /** | **32CV** | **32DC** |
| **EMD code** | EMD-58809 | EMD-58811 |
| **Data collection and processing** |  |  |
| **Magnification** | 150,000 | 150,000 |
| **Voltage (kV)** | 200 | 200 |
| **Electron exposure (e-/Å<sup>2</sup>)** | 40 | 40 |
| **Defocus range (µm)** | -1 to -2.4 | -1 to -2.4 |
| **Pixel size (Å)** | 0.93 | 0.93 |
| **Symmetry imposed** | C1 | C1 |
| **Initial particle images (no.) (from *ab-initio* job)** | 1,666,654 | 2,080,183 |
| **Final particle images (no.)** | 619,311 | 706,064 |
| **Map resolution (Å)** | 3.14 | 2.95 |
| **FSC threshold** | 0.143 | 0.143 |
| **Map resolution range (Å)** | 2.8 to 10.6 | 2.6 to 10.4 |
| **Refinement** |  |  |
| **Initial model used (PDB code)** | 9s4t | 9s2h |
| **Model resolution (Å)** | 3.3 |  |
| **FSC threshold** | 0.5 |  |
| **Map sharpening *B* factors (Å<sup>2</sup>)<sup>a</sup>** | 156 | 131.8 |
| **Model composition** |  |  |
| **Non-hydrogen atoms** | 10886 | 11414 |
| **Protein residues** | 1304 | 1340 |
| **Ligands** | 14 | 22 |
| ***B* factors (Å<sup>2</sup>)** |  |  |
| **Protein** | 69.93 | 56.54 |
| **Ligand** | 88.85 | 76.05 |
| **R.m.s. deviations** |  |  |
| **Bond lengths (Å)** | 0.002 | 0.003 |
| **Bond angles (°)** | 0.546 | 0.620 |
| **Validation** |  |  |
| **MolProbity score** | 1.77 | 2.03 |
| **Clashscore** | 7.48 | 8.34 |
| **Poor rotamers (%)** | 1.60 | 2.25 |
| **Ramachandran plot** |  |  |
| **Favored (%)** | 96.74 | 95.55 |
| **Allowed (%)** | 3.18 | 4.00 |
| **Disallowed (%)** | 0.08 | 0.45 |

## Notes

### Competing Interest Statement

The authors have declared no competing interest.

